# A quantitative framework for interpreting rate and capacity effects in oral regenerative scaffolds

**DOI:** 10.1101/2025.10.25.684540

**Authors:** Maria Teresa Colangelo, Stefano Guizzardi, Carlo Galli

## Abstract

Successful oral and periodontal regeneration requires scaffolds that do more than host cells—they must regulate both the *tempo* and the *extent* of growth within healing tissues. While proliferation data are routinely quantified, comparisons across materials remain fragmented. We introduce the scaffold-augmented logistic (SAL) framework, a minimal quantitative formalism that organizes scaffold performance within a two-parameter space defined by the effective growth rate (*r*_eff_) and the effective carrying capacity (*K*_eff_). Each scaffold can thus be positioned on an R/K design map, where *R* = *r*_eff_/*r*_0_ reflects relative kinetic enhancement and *K*^*^ = *K*_eff_/*K*_0_ expresses sustainable population capacity with respect to a 2D reference. Applied to *in vitro* and *ex vivo* data from oral and periodontal systems, the SAL framework distinguishes R-type (rate-dominated), K-type (capacity-dominated), and hybrid R + K scaffolds. By expressing complex biological outcomes through interpretable parameters, SAL provides a shared quantitative language for scaffold classification and design. It enables model-based reinterpretation of existing data and prediction of design sensitivities, offering a compact quantitative grammar for regenerative scaffold development in oral tissue engineering.

## 1. Introduction

Periodontal and oral tissue regeneration relies on biomaterial scaffolds that provide both structural support and biological guidance for cell colonization and matrix deposition [1]. These scaffolds—ranging from collagen membranes and decellularized matrices to synthetic hydrogels and titanium meshes—serve as temporary extracellular matrices that organize cell attachment, proliferation, and differentiation during healing [2]. Their performance depends not only on biocompatibility but on how they modulate the dynamics of cell growth and survival within a confined three-dimensional environment [3,4]. Successful regeneration therefore requires scaffolds that act as dynamic regulators of population expansion and lineage commitment, rather than passive carriers of cells [5]. These devices function as active microenvironments that can accelerate proliferation, expand sustainable population density, or achieve a balance between the two [6]. Yet, in most studies these effects lack a quantitative framework distinguishing the rate from the extent of cell expansion [7]. The absence of such a shared vocabulary limits cross-study comparison, rational scaffold design, and predictive modeling of biological performance. Mechanistic and computational models have long sought to formalize tissue growth within three-dimensional matrices. Diffusion–reaction frameworks, such as the porous-Fisher equation and multiphase continuum models, successfully capture nutrient gradients, front propagation, and pore bridging dynamics [8]. Agent-based simulations further resolve cell–matrix interactions and local crowding effects [9]. However, these approaches typically emphasize spatial or mechanistic detail rather than reducing scaffold effects to a minimal set of interpretable parameters. Yet, for comparative and design purposes, expressing scaffold influence through just two kinetic quantities—how rapidly cells proliferate and how many can ultimately be sustained—may offer a practical advantage. Although some studies have derived apparent growth constants from proliferation curves [10], these parameters have rarely been used to classify scaffolds systematically.

Scaffolds used in regenerative dentistry encompass a broad spectrum of materials, including polymeric networks, porous foams, decellularized matrices, and bioactive hydrogels with incorporated signaling molecules [11–14]. Comparative evidence consistently shows that architectural and biochemical modifications modulate distinct kinetic dimensions: increased porosity or adhesion sites typically raise the sustainable cell density, while growth-factor incorporation accelerates early proliferation [15]. Despite this recurring pattern, the field still lacks a quantitative framework that integrates both effects within a single formalism capable of linking scaffold architecture to biological performance.

Most models of cell population growth adopt logistic or Gompertz-type equations originally developed for ecological systems [16,17]. The logistic law expresses the interplay between intrinsic proliferation and environmental limitation:

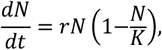

where *N*(*t*) is the number of cells, *r* is the specific growth rate, and *K* is the carrying capacity representing the maximum population sustainable by the environment [18] (Fig. 1). In 2D monolayer culture systems cells experience essentially unlimited access to nutrients and minimal spatial constraint (until confluence) [19], such that the effective growth rate *r* reflects mitotic speed under near-ideal conditions and the carrying capacity *K* corresponds largely to contact inhibition or nutrient depletion at confluence [20,21].

**Figure 1.**
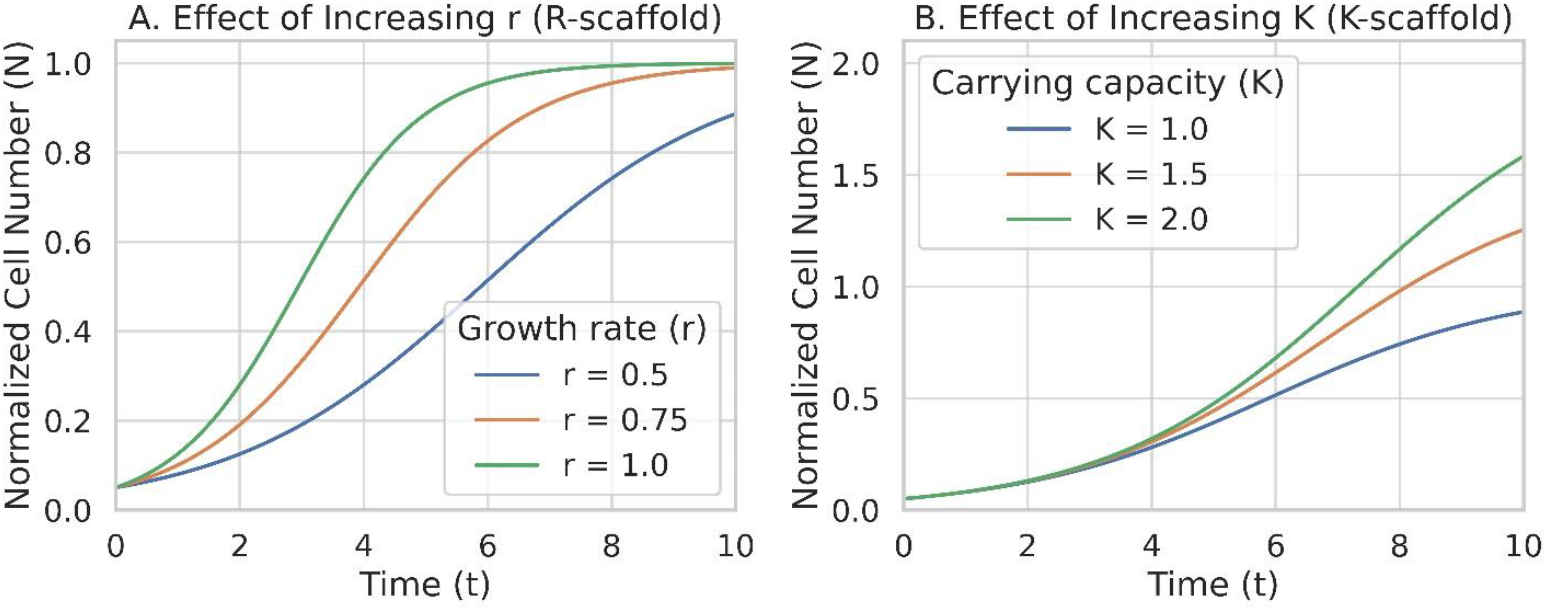
Scaffold-augmented logistic dynamics separate rate from capacity effects. (A) Increasing the effective growth rate *r*_eff_ steepens early proliferation without altering the plateau (R-scaffold behavior). (B) Increasing the effective carrying capacity *K*_eff_ raises the plateau while leaving the early slope largely unchanged (K-scaffold behavior). Parameters are dimensionless and illustrative.

By contrast, in 3D scaffolds both *r* and *K* become emergent properties of the scaffold microenvironment — modulated by pore geometry, nutrient diffusion, and spatial confinement — rather than being intrinsic constants of the cell line [21,22]. Recent theoretical work has extended the logistic family into a continuum of growth laws grounded in resource dissipation and system coherence [23]. In this framework, the classical carrying capacity emerges from feedback between resource exploitation and internal organization, suggesting that even minimal models can capture complex adaptive behaviors. Adapting this logic to 3D scaffolds provides a rationale for reducing their biological influence to two kinetic quantities—rate and capacity—that together describe how efficiently the cellular population exploits the scaffold’s structural and biochemical resources.

Evidence from various in vitro and in vivo studies suggests that modifications in material structure, chemistry, or signaling may differentially influence the apparent proliferation rate and carrying capacity [24]. Porous or adhesive scaffolds tend to support higher final cell densities—consistent with an increase in *K*_eff_ driven by improved nutrient diffusion and surface accessibility [25–27]—whereas growth-factor supplementation or mechanical stimulation typically accelerates early proliferation, indicating a transient rise in *r*_eff_ [28,29]. However, these tendencies are not always disentangled or quantified explicitly. This limitation motivates the present work: to propose a quantitative framework that formalizes how scaffold-dependent microenvironments modulate *r*_*eff*_ and *K*_*eff*_, and to explore whether these parameters can consistently classify experimental outcomes across different materials and cell types. The distinction between rate and capacity is not purely theoretical; it offers a heuristic for interpreting scaffold behavior across applications. For example, materials intended to promote rapid epithelial coverage may benefit from features that transiently enhance *r*_*eff*_, whereas volumetric constructs for bone or connective tissue regeneration may require conditions that sustain a higher *K*_*eff*_. To address this, we introduce the scaffold-augmented logistic (SAL) model, which extends the logistic equation by embedding scaffold-dependent modifiers into *r*_*eff*_ and *K*_*eff*_. This formulation enables a quantitative scaffold classification—the R/K framework—where each scaffold occupies a coordinate in a two-dimensional space defined by relative rate enhancement 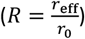 and relative capacity 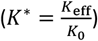. Scaffolds that primarily accelerate proliferation fall along the R-axis (“R-type”), whereas those that expand sustainable cell density lie along the K-axis (“K-type”). Many materials trace intermediate trajectories, such as degradable constructs that initially act as R-type via growth factor release and later transition to K-type as porosity increases.

Because the SAL framework relies on standard proliferation data, it allows both a useful reinterpretation of published experiments and fresh data from new experiments. Fitting *r*_*eff*_ and *K*_*eff*_ across scaffolds yields an empirical R–K map for tissue regeneration (Fig. 2).

**Figure 2.**
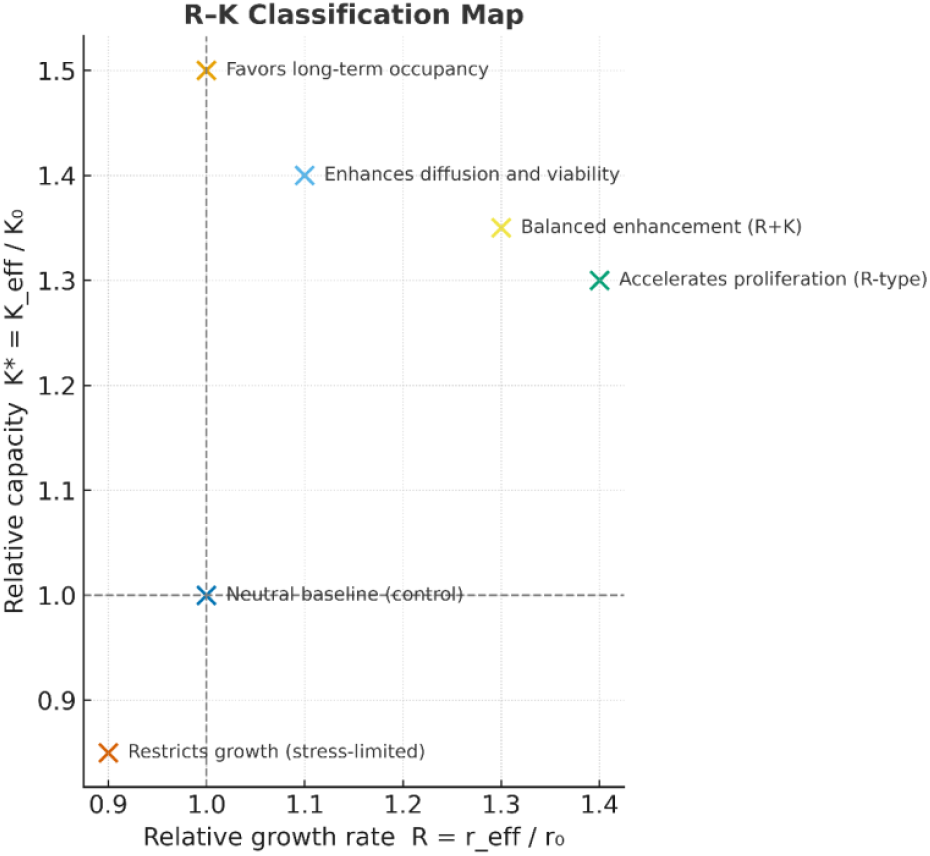
R–K Classification Map. Conceptual representation of scaffold behaviors within the scaffold-augmented logistic (SAL) framework. Each point illustrates a possible combination of relative growth rate and relative carrying capacity compared to a 2D control. Scaffolds that primarily accelerate early proliferation occupy the R-dominant region (right), whereas those that sustain higher long-term occupancy cluster in the K-dominant region (upper). Balanced systems that enhance both parameters lie along the diagonal R+K trajectory. The map provides a visual grammar for comparing how scaffold design modulates kinetic regimes of cell population dynamics.

The main value of the SAL model lies in its parsimony. While multiphysics approaches provide detailed, system-specific insights, the R/K framework offers a complementary abstraction—a simplified but interpretable grammar that allows comparison of scaffold behavior across different studies. The intention is not to replace diffusion–reaction or agent-based models, but to provide a simple unifying layer through which their outcomes can be quantitatively related. The scaffold-augmented logistic framework retains the logistic law as its mechanistic basis, embedding scaffold-dependent modifiers into *r*_*eff*_ and *K*_*eff*_ to highlight how material properties shape measurable kinetics.

## 2. Theoretical Model

The SAL model builds on the logistic law of population growth, but its innovation lies not in mathematical form—it lies in how it translates scaffold effects into two interpretable parameters that can be directly derived from standard proliferation data. Rather than redefining cell dynamics, SAL reframes the logistic equation as a *design map* that connects scaffold descriptors to measurable kinetic outcomes.

We consider cell growth within a three-dimensional scaffold described by a vector of measurable properties

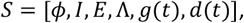

where *ϕ* denotes porosity, *I* interconnectivity, *E* elastic modulus, *Λ* ligand density, *g*(*t*) growth-factor availability, and *d*(*t*) degradation rate. The governing equation becomes

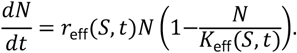

In this formulation, the scaffold acts as an external controller of the intrinsic kinetics. The effective rate term *r*_eff_(*S, t*) captures all dynamic processes that alter cell-cycle frequency, metabolic throughput, or signal-induced proliferation. The effective carrying capacity *K*_eff_(*S, t*) represents the composite capacity imposed by geometry, mechanics, and nutrient accessibility. In degradable or growth-factor-releasing scaffolds, both parameters may evolve over time, reflecting changes in porosity, ligand exposure, or biochemical signaling.

Although both parameters may evolve, particularly in degradable scaffolds or systems with transient growth-factor release, their qualitative influences differ. A positive shift in *r*_eff_ steepens the early slope of the growth curve without changing the ultimate plateau, while a positive shift in *K*_eff_ raises the plateau with minimal effect on the initial kinetics. These features are experimentally distinguishable, allowing scaffolds to be classified empirically by fitting the SAL model to time-course proliferation data.

To illustrate how material properties can be embedded in these parameters, their dependence on scaffold descriptors can be written phenomenologically as:

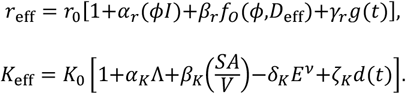

Here, *r*_0_ and *K*_0_ are baseline values from a 2D reference culture; *ϕ* is scaffold porosity; *I* denotes pore interconnectivity; *D*_eff_ the effective diffusivity of oxygen and nutrients; *Λ* the ligand or adhesion-site density; *SA*/*V* the surface area–to–volume ratio; *E* the elastic modulus; and *d*(*t*) a time-dependent degradation term. The functions *f*_0_(*ϕ, D*_eff_) and *g*(*t*) represent, respectively, the effect of oxygen transport efficiency and transient biochemical signaling. The coefficients *α, β, γ, δ, ζ* describe empirical sensitivities of each kinetic parameter to its corresponding property. These relations are not proposed as fitted models but as schematic representations, showing that classical logistic parameters can, in principle, accommodate diverse scaffold-dependent effects within a unified kinetic formalism.

To accommodate spatial heterogeneity, the model can incorporate a diffusion term, leading to a Fisher-type equation:

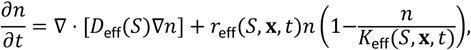

where *n*(*x, t*) represents the local cell density and *D*_eff_(*S*) is the effective diffusivity within the scaffold. This extension allows simulation of pore bridging and spatially nonuniform colonization, phenomena frequently observed in porous collagen and gelatin scaffolds used for gingival fibroblast cultures. Experimental imaging of pore closure shows that higher interconnectivity accelerates front propagation without altering the final occupancy, behavior consistent with an increase in *r*_eff_ but stable *K*_eff_.

From a mechanistic standpoint, *r*_eff_ represents the net outcome of mitogenic stimulation and metabolic availability, while *K*_eff_ reflects the saturation limit imposed by architecture and nutrient supply. This separation enables a compact classification of scaffold behavior into rate-dominated (R), capacity-dominated (K), or hybrid (R + K) regimes.

Beyond classification, the SAL framework offers a foundation for predictive modeling. Regressing experimentally derived *r*_eff_ and *K*_eff_ against scaffold descriptors can reveal design sensitivities—how specific changes in porosity, stiffness, or ligand density shift effective kinetics. Its simplicity is deliberate: the model requires only time-resolved cell number data yet yields parameters that summarize how scaffolds reshape the growth environment. Mathematically minimal, experimentally accessible, and conceptually extensible, the SAL model functions as a unifying scaffold-centric lens for interpreting cell–material dynamics.

## 3. Mechanistic interpretation of scaffold effects on *r*_eff_ and *K*_eff_

Having defined *r*_eff_ and *K*_eff_ as effective descriptors of scaffold-dependent growth, we now examine their biological interpretation. The distinction between *rate* and *capacity* emerges directly from the preceding formulation: changes in *r*_eff_ correspond to processes that accelerate cell-cycle progression or enhance metabolic throughput, while changes in *K*_eff_ reflect modifications in the physical or biochemical limits that determine the maximal sustainable population. In practice, these two dimensions capture complementary aspects of how cells populate a three-dimensional microenvironment—one kinetic, the other structural, and experimental evidence consistently supports this separation. Growth-factor–enriched or otherwise bioactive scaffolds frequently increase *r*_eff_ by activating mitogenic pathways such as MAPK/ERK and PI3K/Akt, leading to earlier metabolic peaks without major changes in final viable-cell density [30]. These systems act as rate enhancers, primarily shifting behavior along the *r*_eff_ axis. Empirical cases confirm these biochemical shifts and reveal dose-dependent trade-offs: Liu et colleagues reported that PLL-grafted electrospun PLGA nanofibers promoted both faster and more extensive proliferation of hUC-MSCs, doubling optical density relative to 2D controls by day 4 and maintaining the advantage through day 8 (*p* < 0.001) [31]. Approximate logistic fits suggest a +30 % increase in *r*_eff_ and an ∼80 % rise in *K*_eff_, consistent with a hybrid R + K response arising from enhanced integrin adhesion and delayed senescence. A biochemical analogue was described by Graziani, who exposed gingival fibroblasts and osteoblasts to graded PRP concentrations [32]. Moderate enrichment (∼2.5× baseline platelets) boosted proliferation by 70–85 %, whereas higher doses suppressed growth and induced differentiation, revealing an R → K transition within a single biochemical continuum. Together, these findings show how trophic signaling can transiently accelerate proliferation before converting to a stabilization-dominant regime.

In contrast, scaffolds with optimized porosity, interconnectivity, or adhesive ligand density—typically achieved through lyophilization, additive manufacturing, or hybrid polymer design—tend to raise *K*_eff_ by improving nutrient diffusion, surface accessibility, and spatial distribution, thus allowing a higher plateau population even when the initial growth rate remains unchanged. Mechanically, substrate stiffness adds another regulatory layer: fibroblasts and osteoblasts sense elasticity through YAP/TAZ-mediated mechanotransduction, with proliferation increasing up to an optimal modulus and declining beyond it [33]. Excessive stiffness restricts deformation and diffusion, reducing long-term viability. In the SAL framework, this produces a non-monotonic dependence of *r*_eff_ and a potential reduction of *K*_eff_ at high crosslinking densities—explaining why some robust hydrogels promote early expansion yet fail to maintain it. Mechanical and fluidic stimuli primarily enhance *r*_eff_. Short-horizon experiments with controlled loading and low-shear dynamics exemplify pure R-type shifts: Nan et al. found that cyclic compression of gingival fibroblasts in 3D PLGA scaffolds (optimal 25 g cm^−2^) increased proliferation and MTT absorbance by ∼60 % after 72 h, corresponding to R ≈ 1.4, *K*^*^ ≈ 1.0—a pure r-type modulation mediated by TGF-β1 signaling [34]. Similarly, hDPSCs cultured under simulated microgravity exhibited higher Ki-67 positivity and faster ECM network formation than static controls, indicating a dynamic-culture-driven r-boost that seeded later differentiation [35]. These results illustrate the mechanical–fluidic corner of the R-dominant quadrant, where transient kinetic acceleration precedes structural stabilization.

Porosity and interconnectivity also control *K*_eff_ indirectly through mass-transport limitations. In diffusion-restricted constructs, oxygen and nutrient depletion create internal gradients that suppress proliferation and induce quiescence or necrosis, effectively lowering the apparent carrying capacity even if *r*_eff_ remains intrinsically high. Imaging and oxygen-mapping studies confirm that scaffolds with pores above roughly 200 µm sustain larger viable fractions than those below 100 µm, which often develop hypoxic cores. Within the SAL model, these observations correspond to spatially varying *r*_eff_(*x, t*) and a global reduction in *K*_eff_, consistent with porous-Fisher predictions. These mechanisms are mirrored quantitatively in recent 3D systems: Zenobi and colleagues cultured osteoblast-like cells (SAOS-2, U2OS) on 3D-printed PLA scaffolds mimicking either healthy or osteoporotic trabecular bone [36]. Under dynamic rotary culture, the osteoporotic design supported nearly a threefold increase in viable cell density over static controls after 14 days. Early proliferation rates were comparable, indicating that the enhancement derived from improved *K*_eff_, not *r*_eff_. Approximate logistic fitting (R ≈ 1.0, *K*^*^ ≈ 3.0) identifies a clear K-type response driven by superior mass transport and accessible surface area. The same capacity-dominated pattern was confirmed in trabecular titanium scaffolds fabricated via Electron-Beam or Selective-Laser Melting. Human adipose-derived MSCs proliferated steadily on both architectures, reaching higher late-time plateaus (R ≈ 1.0, *K*^*^ ≈ 1.6–1.8) with minimal change in early slope. Their large pores (∼640 µm, 65 % porosity) enabled deep infiltration and long-term viability. Increased expression of COL1A1, OSX, and mineralized ECM confirmed that 3D structure itself acts as a capacity amplifier—enhancing *K*_eff_ through geometric and mechanical optimization while leaving *r*_eff_ stable [37].

A related mechanism was observed by Peluso, who combined a synthetic PCL backbone with a hydrophilic chitosan network. PDLSCs and MG-63 cells cultured on this dual-porosity composite displayed ∼20 % higher viability at day 14 compared with pure PCL scaffolds. Logistic refitting (R ≈ 1.02, *K*^*^ ≈ 1.20) identified a mild but consistent K-type shift, attributable to improved water uptake and cell–matrix contact [38]. Collectively, these three studies outline the architectural K-axis of the SAL map: improved diffusion and surface accessibility increase *K*_eff_ and sustain larger steady-state populations without accelerating division.

Compositional tuning produces analogous capacity-dominant effects: osteoblasts cultured on collagen–GAG scaffolds exhibited faster proliferation on collagen-rich, stiffer matrices (r-shift) and slower but more stable growth on GAG-enriched ones (K-shift) [39]. Similarly, Shor et al. showed that adding hydroxyapatite (HA) to PCL scaffolds did not change proliferation kinetics but enhanced ALP activity and mineralized matrix deposition, indicating elevated *K*_eff_ at constant *r*_eff_ [40]. Both systems exemplify K-type modulation through biochemical or mineral reinforcement—increasing the sustainable population capacity by stabilizing the tissue-like phase rather than by speeding mitosis.

In periodontal models, the same logic holds: PDGF-enriched collagen predominantly shifts r, macroporous or decellularized matrices elevate K, and fibrin constructs show slower onset but higher eventual occupancy.

Gingival fibroblasts cultured in collagen scaffolds supplemented with platelet-derived growth factors display sharply increased metabolic activity within 48 hours, yet their final density matches that of unloaded controls. Fitting these curves with the SAL equation yields a 1.5–2.0-fold increase in *r*_eff_ with stable *K*_eff_—a signature of R-type behavior. Beyond their mitogenic role, many growth factors such as PDGF and FGF also exert potent chemotactic effects, promoting directed cell migration into the scaffold rather than simply accelerating division [41]. In the SAL framework, this dual influence would appear as a transient local increase in *r*_eff_(*x, t*) at the colonization front and, over time, as an effective rise in *K*_eff_ as the newly recruited cells expand the occupied volume. This behavior aligns with the Fisher-type spatial extension of the model, where diffusion-driven infiltration and proliferation are coupled. Consequently, growth-factor-enriched scaffolds should not be interpreted solely as R-type accelerators: their chemotactic signaling can indirectly enhance capacity by facilitating more complete spatial occupancy. Conversely, fibroblasts seeded in decellularized gingival matrices or highly porous collagen sponges exhibit similar initial kinetics but reach higher DNA content at later time points, corresponding to a 1.3–1.5-fold rise in *K*_eff_ with minimal change in *r*_eff_. Osteoblast cultures on hydroxyapatite–collagen composites show analogous trends: increasing porosity raises the plateau population (higher *K*_eff_) without affecting early slopes, while addition of BMP-2 selectively boosts *r*_eff_. Such analyses suggest that what once appeared as heterogeneous outcomes can be organized coherently along the R–K axes. From the proliferation curves reported by Agis and colleagues for human gingival fibroblasts cultured in collagen scaffolds, approximate fits yield *R* ≈ 1.4 and *K*^*^ ≈ 1.4 for the sponge + PDGF condition, and *R* ≈ 1.3, *K*^*^ ≈ 1.1 for the gel + PDGF condition, relative to the baseline gel scaffold (*R* = *K*^*^ = 1) [42]. These estimates place the PDGF-loaded sponge scaffold in the upper-right quadrant of the R–K plane—enhancing both proliferation rate and sustainable population density—while the PDGF-loaded gel aligns closer to the R-axis, reflecting a primarily rate-driven effect. Similarly, data from Ardakani et al. suggest a modest capacity uplift (*K*^*^ ≈ 1.2) in a collagen membrane variant, with limited change in early growth (*R* ≈ 1.1) [43]. In contrast, proliferation patterns of gingival fibroblasts cultured in 3D fibrin scaffolds indicate a K-dominated response, with slower early kinetics but higher final occupancy (*R* < 1, *K*^*^ > 1) [44].

To further illustrate the practical extractability of SAL parameters, we re-fitted unpublished proliferation data for human gingival fibroblasts cultured on plastic in the absence or in the presence of hyaluronic acid (HA). Logistic fitting yielded baseline parameters of r_0_ ≈ 0.055 h^−1^ and K_0_ ≈ 2.8 × 10^5^ cells, versus *r*_eff_ ≈ 0.057 h^−1^ and *K*_eff_ ≈ 3.1 × 10^5^ cells for HA-treated cultures. These correspond to an approximately +4 % increase in the effective growth rate and a +10 % increase in the carrying capacity. The disproportionate gain in K relative to r indicates that HA primarily enhances the sustainable population size rather than accelerating early proliferation. Within the SAL framework, this behavior identifies HA as a capacity-oriented (K-type) scaffold that favors long-term tissue stability over short-term proliferative boost.

At micrometer scale, patterned implant surfaces act as quasi-3D scaffolds that bias r and K through mechanotransduction and surface-energy cues: Rabel showed that anisotropic titanium and zirconia surfaces with moderate roughness increased the fraction of dividing osteoblasts (high *r*_eff_), whereas isotropic, highly enlarged surfaces reduced proliferation but supported greater matrix deposition (high *K*_eff_) [45]. This geometry-encoded R↔K transition was mirrored by Schweikl and Andrukhov, who demonstrated that increasing titanium roughness suppressed early fibroblast proliferation but enhanced matrix synthesis, cytokine signaling, and oxidative stress adaptation—traits of K-type stabilization [46,47].

Scaffold degradation further introduces temporal dynamics: resorbable or swelling materials progressively modify diffusivity, stiffness, and surface exposure, effectively tracing a trajectory through the R–K plane. Early in culture, soluble cues or surface charge favor faster proliferation, increasing *r*_eff_; as degradation advances and porosity rises, *K*_eff_ grows, permitting continued expansion. This temporal evolution explains the biphasic kinetics frequently observed in degradable hydrogels and polymer composites—an initial burst of proliferation followed by a second growth phase driven by architectural remodeling.

Together, these mechanisms outline a simple quantitative logic: biochemical and mechanical stimuli move a scaffold rightward along the rate axis by accelerating proliferation, whereas improvements in geometry, surface area, or diffusion shift it upward along the capacity axis by expanding sustainable occupancy. Multifunctional systems that combine these effects trace diagonal or time-dependent paths, achieving both rapid colonization and long-term stability. Conversely, antagonistic factors—such as excessive stiffness coupled with poor diffusion—can yield transient rate enhancement followed by premature saturation. In this way, the SAL framework translates diverse mechanobiological behaviors into trajectories on a defined two-parameter plane rather than separate descriptive categories.

This mapping also generates explicit, testable predictions. A porous membrane incorporating controlled mitogen release should exhibit a combined rightward-and-upward displacement relative to an unloaded dense control, indicating concurrent enhancement of *r*_eff_ and *K*_eff_. A highly crosslinked hydrogel with limited permeability would occupy a region of elevated *r*_eff_ but reduced *K*_eff_, corresponding to fast early proliferation followed by stagnation. Such hypotheses can be directly verified by fitting experimental N(t) curves to the SAL model and comparing normalized parameters *r*_eff_/r_0_ and *K*_eff_/K_0_. Degradation and remodeling then transform the static map into a dynamic trajectory describing the scaffold’s life cycle.

In vivo, the same logic extends beyond cell culture. Once implanted, *K*_eff_ becomes influenced by vascularization, immune crosstalk, and host integration. Materials that function as strong R-boosters in vitro may fail clinically if they cannot sustain *K*_eff_ post-implantation, while high-capacity but slow-proliferating scaffolds may integrate poorly due to limited early colonization (Fig. 3). Locating where a scaffold sits—or how it moves—within the R–K plane therefore provides a practical design compass: R-dominated features favor rapid closure and cell recruitment, K-dominated features sustain long-term tissue consolidation.

**Figure 3.**
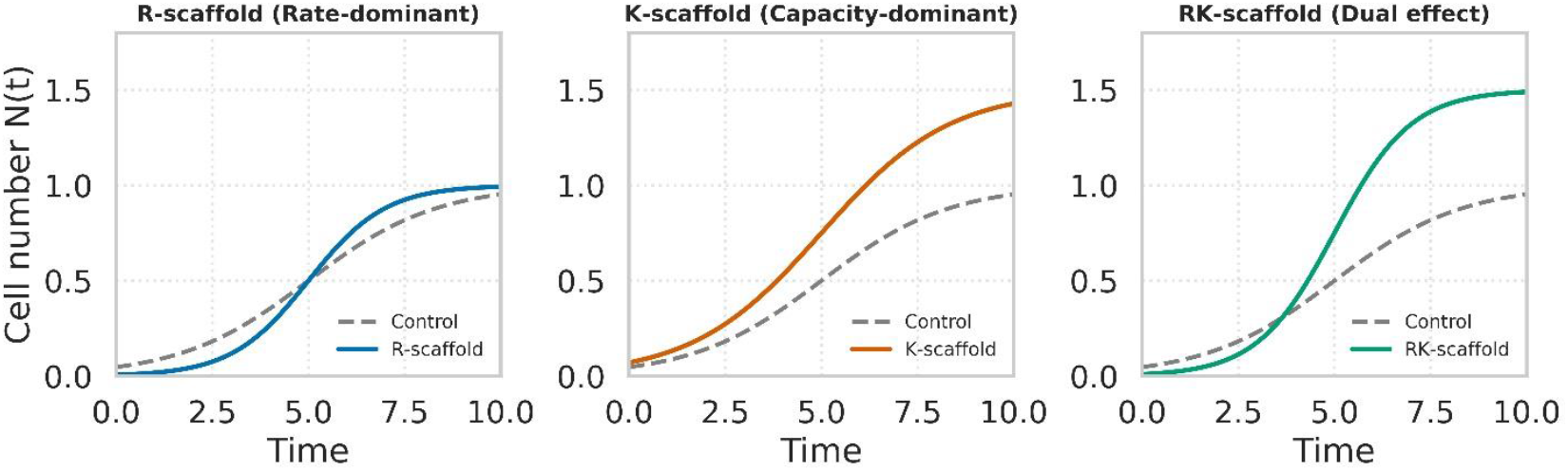
Representative growth dynamics for R-, K-, and RK-scaffolds. R-scaffolds (blue) increase the growth rate *r*_*eff*_ without altering the carrying capacity *K*_*eff*_ K-scaffolds (orange) raise *K*_*eff*_ without accelerating early proliferation; RK-scaffolds (green) enhance both parameters, combining rapid colonization with higher sustainable cell density. Dashed gray lines represent control behavior.

## 4. Discussion: Integrating the SAL framework with experimental data and scaffold design

The conceptual strength of the scaffold-augmented logistic (SAL) framework lies in its ability to reinterpret a vast and fragmented experimental literature through a unified quantitative language [27]. Most studies on tissue scaffolds report time-dependent proliferation curves—often fitted with exponential or polynomial functions that yield parameters with limited biological interpretability [48]. Recasting those same data within the SAL formalism produces two interpretable quantities, *r*_eff_ and *K*_eff_, which immediately reveal whether a material enhances the *tempo* or the *extent* of cellular expansion. This simple translation can transform descriptive growth data into design-relevant information.

This framework also opens a path to predictive design. Because *r*_eff_ and *K*_eff_ can be expressed as functions of measurable scaffold descriptors, empirical or semi-empirical relations can link microstructural parameters to biological outcomes [49]. A regression analysis across studies could quantify, for instance, how a 10 % increase in porosity or ligand density translates into proportional changes in *r*_eff_ or *K*_eff_ [50]. Once such sensitivities are known, scaffold optimization becomes a more tractable engineering problem rather than an empirical exercise. In principle, the same model could drive computational searches for scaffold architectures that achieve prescribed positions within the R–K plane—fast-acting for epithelial closure, or capacity-dominant for connective-tissue augmentation. Beyond parameter optimization, the same quantitative framework can be embedded within spatial or mechanistic models to simulate how those optimized parameters manifest in real 3D geometries.

The SAL formalism remains compatible with more detailed spatial models [51]. The Fisher-type extension naturally connects to diffusion-limited proliferation and pore-bridging kinetics, enabling simulation of colonization dynamics in complex geometries such as interdental defects. When local *r*_eff_(*x*) and *K*_eff_(*x*) are inferred from scaffold architecture, the model predicts macroscopic quantities such as time to full pore occupation—metrics that experimentalists already use as surrogates of performance. In this sense, the SAL model acts as the missing intermediary between the simplicity of logistic fitting and the complexity of multiphase or agent-based simulations.

Like any reduced model, the SAL framework abstracts biological detail. It assumes partial independence between *r*_eff_ and *K*_eff_, though in practice scaffold properties often influence both. For example, increasing porosity enhances diffusion (raising *r*_eff_) while simultaneously enlarging available space (raising *K*_eff_); degradation processes can likewise affect both terms at once. The framework captures the resultant behavior but cannot disentangle these couplings without factorial experimental designs that vary single parameters systematically. A second limitation concerns the interpretation of *K*_eff_, which, although mathematically an asymptotic population size, integrates multiple constraints—geometric, metabolic, and mechanical. Refining its biological meaning requires complementary measurements such as oxygen mapping or matrix deposition assays.

Reliable estimation of *r*_eff_ and *K*_eff_ requires well-designed growth assays with sufficient temporal resolution and independent measures of cell number. While many legacy studies do not meet these standards, new experiments can be explicitly structured for SAL fitting—using multiple seeding densities, direct DNA or nuclei quantification, and extended observation to saturation. Under such conditions, *r*_eff_ and *K*_eff_ become experimentally separable and reproducible, enabling systematic exploration of how scaffold properties drive kinetic shifts. Oxygen and substrate depletion, as well as changes in mitochondrial efficiency, can distort endpoint readouts even when cell numbers remain constant. Therefore, metabolic dyes such as MTT, resazurin, or alamarBlue should be interpreted primarily as indicators of early proliferative activity—that is, as proxies for *r*_eff_—rather than as accurate reflections of the carrying capacity [52]. Quantitative estimation of *K*_eff_ should rely on absolute counts, DNA quantification, or nuclei imaging whenever possible [53]. Under these circumstances, the fitted values of *r* and *K* can become correlated and non-unique, making it impossible to ascribe changes reliably to one parameter or the other.

We fully acknowledge that scaffold features such as porosity or stiffness seldom act on only one of the parameters *r*_eff_ or *K*_eff_. In practice, porosity can improve diffusion (raising *r*_eff_) and simultaneously enlarge available space or adhesion surface (raising *K*_eff_). Similarly, changes in stiffness may alter mechanotransduction signals that boost proliferation [54] and at the same time mechanically constrain the maximum occupancy or alter degradation dynamics that ultimately modify the accessible space. These effects are often coupled rather than orthogonal. For instance, increasing porosity may initially accelerate nutrient transport and early proliferation but later shift the equilibrium toward higher saturation density—a trajectory that simultaneously elevates both *R* = *r*_eff_/*r*_0_ and *K*^*^ = *K*_eff_/*K*_0_. Hybrid hydrogels combining biochemical cues with structural modifications often exhibit this behavior, showing an early kinetic boost followed by enhanced long-term occupancy.

By contrast, materials engineered to isolate one mechanism provide useful boundary cases. Unidirectional collagen matrices with high swelling capacity but unchanged proliferation kinetics [55] predominantly affect *K*_eff_ with minimal change in *r*_eff_. Conversely, fibroblast cultures supplemented with basic FGF (10–100 ng mL^−1^) display steeper early growth curves and unchanged final plateaus [56] consistent with a selective increase in *r*_eff_ alone. These empirical observations define the two limiting cases—pure K-type and pure R-type—within a broader continuum where most scaffolds exert mixed or evolving influences.

When multiple scaffold parameters vary simultaneously—such as porosity with ligand density or stiffness with degradation rate—the resulting shifts in *R* and *K*^*^ reflect combined effects that cannot be uniquely decomposed. Disentangling these contributions would require factorial designs where individual parameters are independently varied and monitored across multiple seeding densities and time points. Until such datasets become available, the SAL framework is best applied as a quantitative descriptor of *net kinetic shifts*, rather than as a tool for inferring mechanistic causation. Ultimately, the SAL model provides a practical and conceptually transparent bridge between data and design. It converts routine proliferation measurements into parameters that carry mechanistic meaning, establishing a vocabulary shared across disciplines (Fig. 4). By reformulating scaffold behavior in terms of rate and capacity, the SAL framework offers a quantitative interface between materials science and cell biology, providing a common reference space for relating scaffold architecture to biological response.

**Figure 4.**
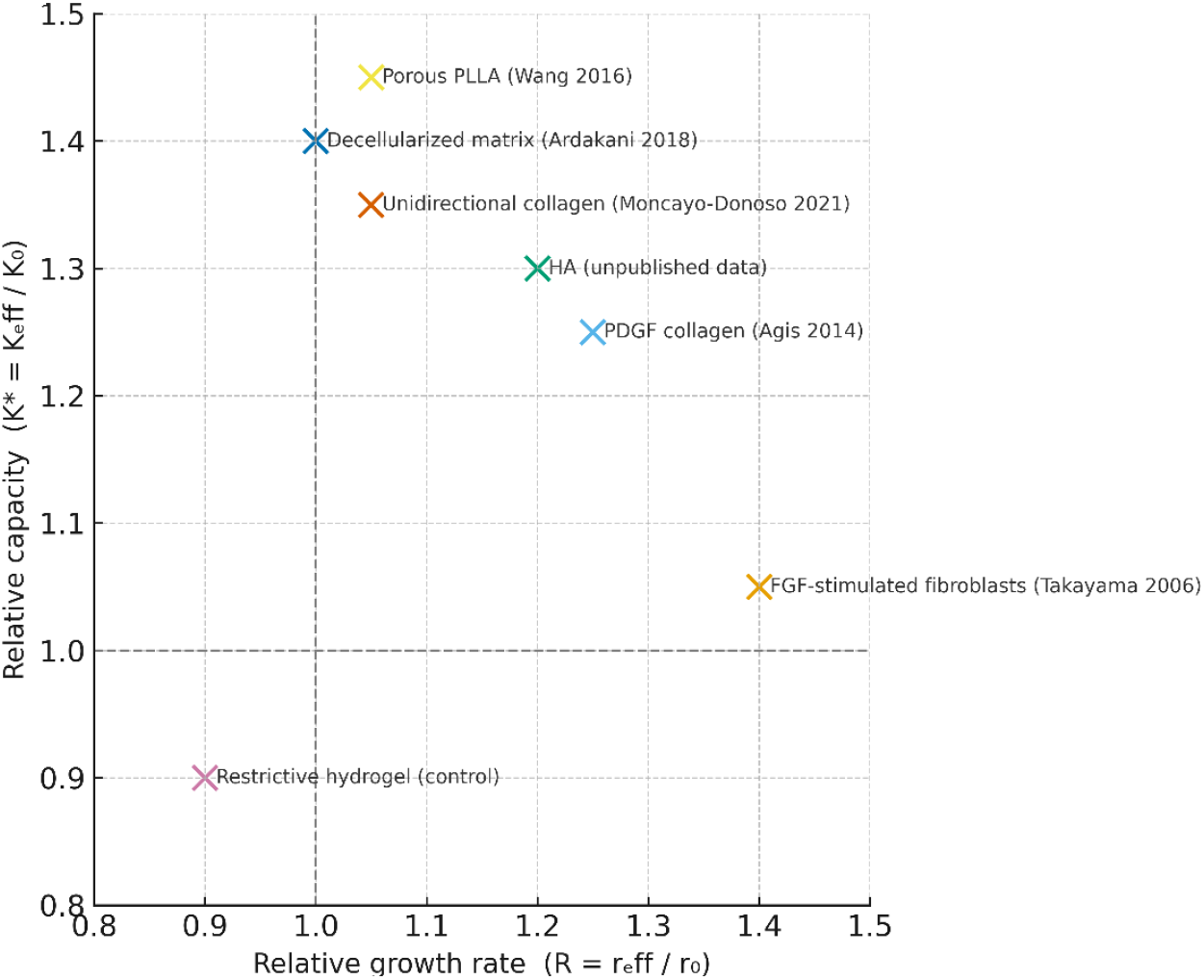
R–K Classification Map of Scaffold Behaviors. Each point represents a scaffold system positioned according to its relative proliferation rate and carrying capacity compared to 2D culture. Data are drawn from representative studies: FGF-stimulated periodontal fibroblasts (Takayama 2006), PDGF-loaded collagen membranes (Agis 2014), porous PLLA scaffolds (Wang 2016), decellularized gingival matrices (Ardakani 2018), and unidirectional collagen matrices (Moncayo-Donoso 2021), together with Hyaluronic acid hydrogel experiments (unpublished data). Rightward displacements indicate enhanced proliferative rate, upward displacements greater sustainable occupancy, and diagonal shifts balanced modulation of both parameters.

## 5. Implications and future directions

The scaffold-augmented logistic framework reframes a central question in oral and periodontal regeneration [11]. Instead of asking whether a scaffold enhances cell colonization, it asks *how*—by accelerating the growth rate or by increasing the sustainable population it can support. The distinction is not between qualitative and quantitative analysis, but between isolated quantitative metrics and a unified parametric representation that enables comparison across materials and studies. This shift has both theoretical and translational implications, providing a common coordinate system that links scaffold design parameters to biological performance.

For early wound healing, where rapid epithelial closure and fibroblast expansion are decisive, R-type scaffolds provide the functional advantage [57]. Materials that release growth factors or present compliant, cell-permissive mechanics act as kinetic amplifiers, shortening the proliferative phase and reducing time to defect coverage [49]. In contrast, long-term regenerative goals—such as stable connective-tissue augmentation or bone fill—depend on K-type behavior, in which the scaffold sustains large, viable populations over extended periods through robust architecture and efficient diffusion. Recognizing that these functions may compete allows rational design trade-offs rather than empirical compromise.

The immediate next step is data integration. Thousands of published studies already contain the necessary inputs—cell number or metabolic-activity curves over time—but few extract quantitative growth parameters [58]. A systematic meta-analysis applying the SAL model could generate distributions of *r*_eff_ and *K*_eff_ across material classes and correlate them with descriptors such as porosity, stiffness, or ligand density. The resulting R–K map of the biomaterial landscape would highlight unexplored regions: materials that balance both parameters or exhibit time-dependent transitions between them. Such mapping could help transform isolated experiments into a collective dataset for hypothesis-driven scaffold design.

Temporal extensions of the model add further relevance. In degradable scaffolds, *K*_eff_(*t*) typically increases as structure opens, while *r*_eff_(*t*) may decline as growth-factor reservoirs deplete. Modeling these coupled trajectories can predict population dynamics during healing and link scaffold degradation with tissue remodeling. This dynamic interpretation transforms the R/K plane from a static classification into a kinetic landscape describing the scaffold’s life cycle.

The SAL framework also forms a natural interface with data-driven and machine-learning approaches. Once sufficient *r*_eff_ and *K*_eff_ values are available, algorithms can learn the nonlinear mapping from design parameters to biological outcomes, allowing prediction of scaffold behavior without exhaustive experimentation. However, realizing this potential will require more than standardized reporting of growth curves: it demands reproducible kinetic assays, consistent normalization procedures, and transparency in data fitting. Establishing such quantitative discipline across laboratories remains a significant but necessary step toward making scaffold design genuinely predictive.

In conclusion, the scaffold-augmented logistic model provides a minimal yet versatile framework for relating material design to biological performance. By distinguishing between rate and capacity effects, it offers a quantitative language for interpreting how scaffold properties shape cell growth dynamics. As scaffold technologies become more complex and multifunctional, such integrative approaches may prove increasingly valuable for linking experimental observation to predictive understanding in regenerative research.

